# Catalytic relevance of quinol anion in biological energy conversion by respiratory complex I

**DOI:** 10.1101/2024.09.06.611712

**Authors:** Oleksii Zdorevskyi, Johannes Laukkanen, Vivek Sharma

## Abstract

Redox chemistry of quinones is an essential component of life on earth. In the mitochondrial electron transport chain, ubiquinone molecule is reduced to ubiquinol by respiratory complex I to drive the synthesis of ATP. By performing both classical and hybrid QM/MM simulations on high-resolution cryo-EM structures, including quantitative free energy calculations, we show that semiquinone species in complex I is anionic in nature and is trapped in the active site chamber for its subsequent reduction. Two-electron reduction of ubiquinone yields a metastable ubiquinol anion, which is electrostatically pushed by 15-20 Å towards the exit of the ubiquinone binding chamber to drive the proton pump of complex I. As part of the two-electron reduction of ubiquinone, protonic rearrangements take place in the active site in which a highly conserved histidine converts from its one tautomeric state to another. The combined findings provide a detailed and testable mechanistic picture of proton-coupled electron transfer reaction at the active site of complex I in wild-type as well as mutant conditions.

## Introduction

Quinone molecules are redox-active elements found in all domains of life. By constantly shuttling between their reduced and oxidized forms, they facilitate the transfer of electrons (and protons) to and from various energy converting enzymes [1]. In the inner membrane of mitochondria, which harbours the electron transport chain, long-tailed ubiquinone (UQ) molecules transfer electrons between membrane-bound redox-active proton pumping enzymes. The first enzyme of this chain, respiratory complex I (Fig. 1), is a NADH:ubiquinone oxidoreductase, and couples the oxidation of NADH (E_m,7_ ∼ –320 mV) with the two-electron reduction of UQ (E_m,7_ ∼ +100 mV). The energy released during UQ reduction is employed to pump protons across the inner mitochondrial membrane [2,3] (Fig. 1), creating a proton electrochemical gradient, which drives ATP synthesis and active transport [4].

**Fig. 1.**
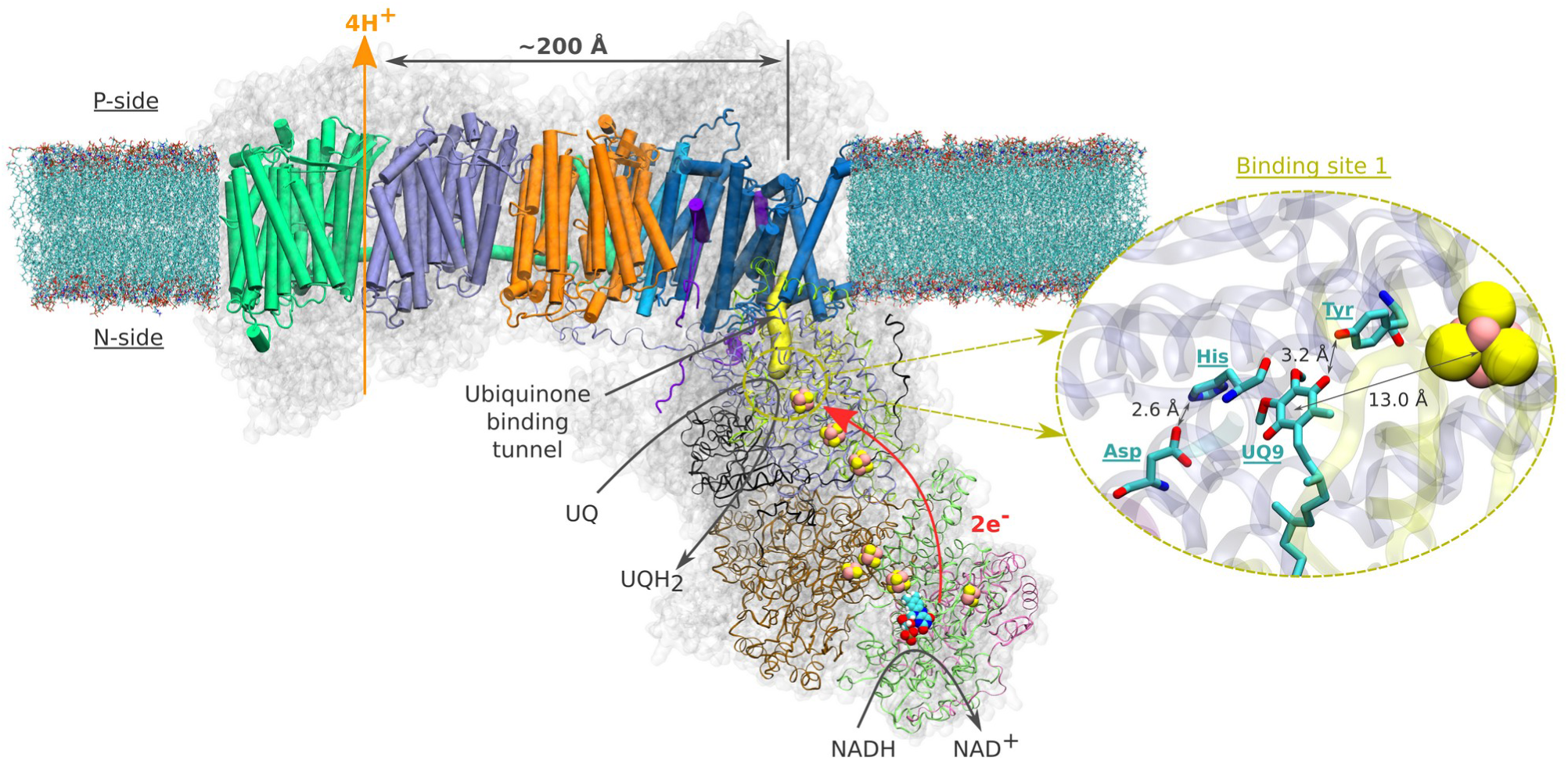
Structural organisation of the mitochondrial respiratory complex I from *Yarrowia lipolytica* [27] (PDB 7O71). Membrane-bound core subunits are shown in cartoon representation, whereas the core subunits of the peripheral arm in ribbon representation. The accessory subunits are represented with a transparent grey surface. Van der Waals representation is used for the redox cofactors of the peripheral arm (iron sulfur clusters and FMN molecule). Yellow surface represents the UQ chamber – a protein cavity providing access for the UQ species to the redox site (plotted with CAVER (v. 3.0) [36], using ^49kDa^His95 and ^49kDa^Tyr144 as starting points, and a 0.7 Å cutoff). The inset illustrates the binding of UQ at site 1 [9]. Spatial orientation of UQ head group and the adjacent titratable residues are based on the structure [16] from *Sus scrofa* (PDB ID: 7V2C).

Despite an extraordinary amount of high-resolution structural data on complex I for over a decade, its redox-driven proton pumping mechanism still remains elusive and extensively debated [5–8] (Fig. 1). According to the current consensus (see various proton pumping models discussed in ref [6]), a UQ molecule enters from the membrane milieu to the ∼30 Å long UQ binding chamber (UQ tunnel) and extracts electrons from the terminal FeS cluster N2 (Fig. 1). During tunnel travel, UQ binds to several intervening binding sites (sites 1-5) within the UQ tunnel that were predicted by molecular dynamics simulations [9–12] and structurally characterized by cryo-EM [13,14]. At the deep binding site 1, UQ head group binds in the vicinity of a conserved His/Asp pair (^49kDa^His95/^49kDa^Asp196 – *Yarrowia lipolytica* complex I numbering) and stabilised by a hydrogen bond with conserved tyrosine (^49kDa^Tyr144) sidechain (Fig 1, inset) [6,15–19]. Bound within the electron transfer distance (<15 Å) from the N2 FeS center (see Fig. 1), reduction of UQ at site 1 drives electrostatic and conformational transitions [3,6,15,18,20] that are thought to be important for the coupling mechanism. However, the exact sequence of charge transfer and conformational events remains unknown, resulting in incomplete understanding of redox-driven proton pumping mechanism of complex I. Here, we performed classical and hybrid QM/MM (quantum mechanical/molecular mechanical) MD simulations on high-resolution cryo-EM structures of complex I to study the protonation and tunnel dynamics of two species of UQ (ubisemiquinone and ubiquinol) that are formed after one- and two-electron transfer, respectively. Our data suggests that the protonation of ubisemiquinone (USQ) is energetically unfavorable, and posits anionic ubiquinol (UQH^-^) as a catalytically competent species in the redox catalysis of complex I. We outline a step-by-step mechanistic picture of the UQ reduction and dynamics and provide molecular basis to previously enigmatic site-directed mutagenesis data.

## Results

### Energetics of protonation and dynamics of one electron reduced UQ species

Electrons are transferred from the terminal N2 cluster to bound UQ one at a time, leading to an inevitable formation of ubisemiquinone (USQ) radical. Our hybrid (QM/MM) free energy simulations on the high-resolution structure of respiratory complex I (see methods) show that the protonation of USQ by nearby residues is unfavorable by 8-10 kcal/mol (Fig. 2A). Spin density analysis reveals that the electron is indeed localized on the UQ headgroup (Fig. S1A), however, a proton transfer does not occur to it, neither from the hydrogen bonded tyrosine nor from the His/Asp pair. At the same time, noticeable protonation and conformational rearrangements take place during QM/MM minimization and simulations. The histidine residue, which is initially modelled as doubly protonated (imidazolium), donates its proton to the anionic aspartate in both the fully oxidized and one-electron reduced states, thereby rendering both the residues charge-neutral (Fig. S2A). This is in agreement with the classical free energy simulation data that showed charge-neutral states of histidine and aspartate [17]. Interestingly, our simulation data revealed that histidine can drift from its structural position (as in PDB 7V2C [16]) to form a stable hydrogen bond with the ketonic group of UQ within the timescales of QM/MM simulations (Movie. S1). This conformational rearrangement brings His/Asp system in a pose ready for proton donation upon second electron transfer (see below).

**Fig. 2.**
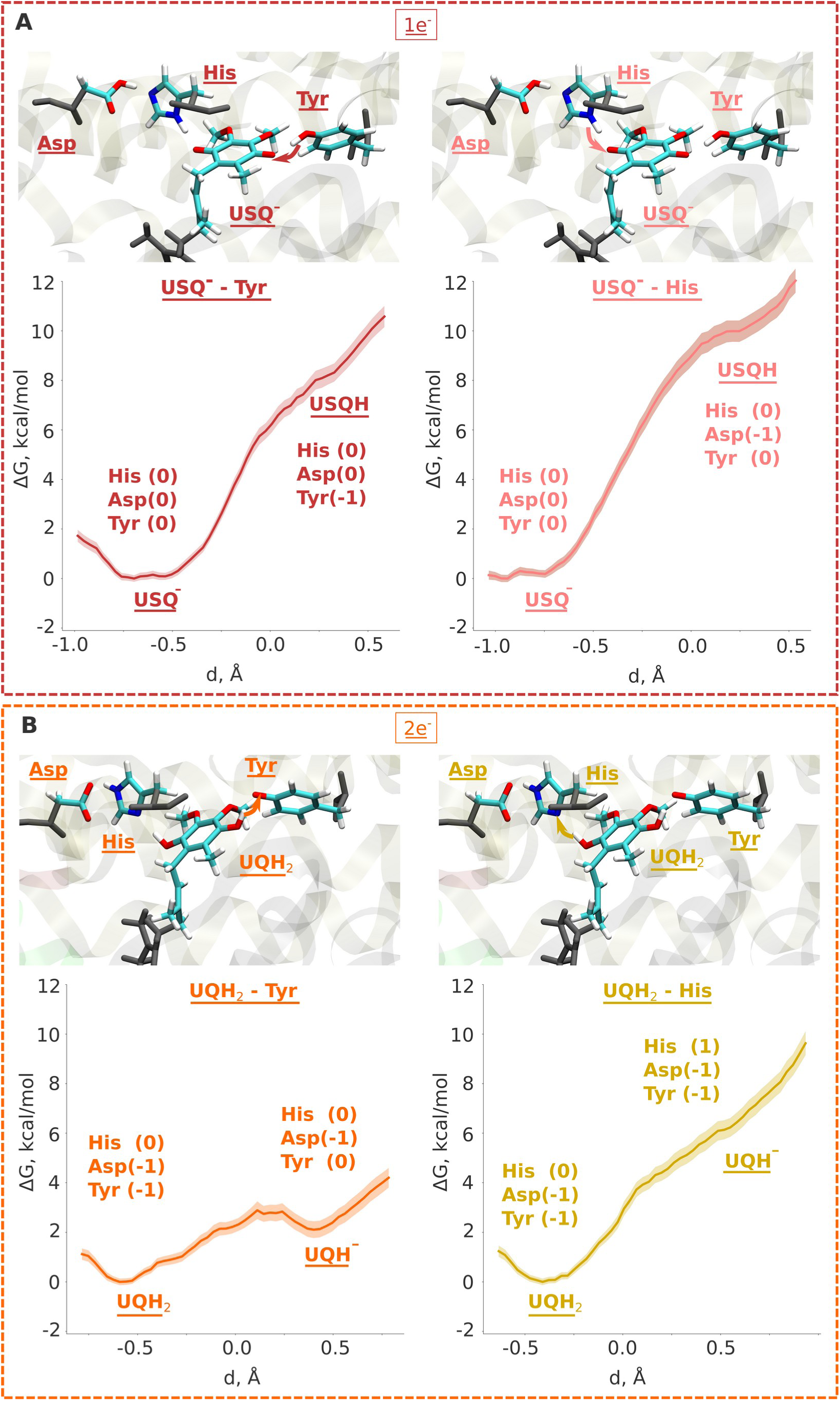
QM/MM umbrella sampling free energy profiles of the protonation dynamics of ubiquinone in one-(**A**) and two-electron (**B**) reduced cases. (**A**) Proton transfer from Tyr (left panel), and from His (right panel) to anionic ubisemiquinone USQ^-^. (**B**) Proton transfer from Tyr (left panel), and from His (right panel) to anionic ubiquinol UQH^-^. Charge states of the protein residues are shown as a caption to each state. The upper insets show simulation snapshots (taken from unbiased MD simulations) with arrows pointing out proton transfer simulated by free energy simulations (see methods). The backbone atoms from the MM region are coloured in grey. The whole QM region is shown in Fig. S10.

Even though transient formation of USQ radical is inevitable, analysis of EPR [21,22] have raised concerns about the origins and occupancy of USQ species [6]. The long-life of a free radical state like USQ and its uncontrolled diffusion can lead to proteolipid oxidation and formation of dangerous ROS [23]. Previous non-equilibrium classical MD simulations showed that a higher amount of force (work) is required to *pull* a charged USQ species out of the UQ tunnel, leading to the suggestion that keeping USQ anionic is a strategy to trap it in the UQ tunnel [10]. To probe the long-time scale dynamics of anionic USQ, we performed microsecond-long unbiased equilibrium MD simulations (see methods). In contrast to the stable position of oxidised UQ near the N2 FeS center (Fig. 3), the singly reduced UQ (USQ^-^) showed partial unbinding and diffusion towards the UQ binding site 4, closer to the entrance of the UQ tunnel. The higher stability of oxidized UQ compared to USQ^-^ is in part due to the stable hydrogen-bonding interactions with the neighbouring residues (Fig. S3). Because unbiased MD simulations are not able to sample the tunnel dynamics of UQ and also to probe the dynamics in a quantitative manner, we performed classical US-based free energy simulations (see methods). We find that even though USQ^-^ is in part unstable at site 1 with respect to oxidized UQ (Fig. 4), its diffusion towards the tunnel exit is also restricted by a similar ∼4 kcal/mol energy barrier. This barrier is in part caused by the frequent hydrogen bonds of the USQ^-^ head group with the histidine (Fig. S3B). This will not only lead to the trapping of USQ^-^ at the site near N2 but will also prime it for fast reduction by a second electron transfer, most likely within the nanoseconds of first electron transfer.

**Fig. 3.**
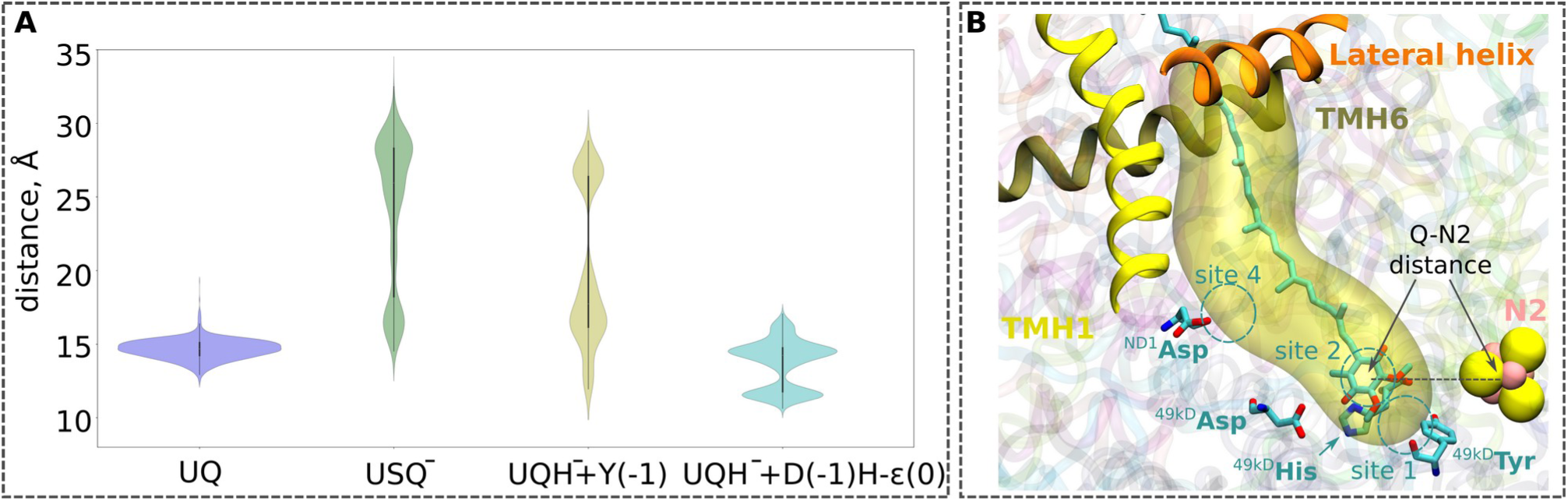
A. Violin plots for the UQ-N2 distances from the MD simulations (1 μs x 3 replicas) of high-resolution structure of respiratory complex I from *Yarrowia lipolytica* (PDB ID: 7O71) [27] with quinone modelled at site 1 (see panel **B**). The data for UQ are shown in blue, for USQ^-^ in green, and for UQH^-^ in yellow. For the UQH^-^ dataset, ^49kDa^Tyr was additionally deprotonated to model the proton transfer from Tyr to the double-reduced quinone (yellow plot). The second UQH^-^ dataset was collected from simulations with an ε-state of ^49kDa^His, and deprotonated ^49kDa^Asp (cyan plot), modelling proton donation from the His/Asp pair to doubly-reduced UQ. The time series plots are shown in Fig. S12. **B** Illustration of the distance between the N2 iron-sulfur cluster and the ubiquinone head group (UQ-N2 distance) plotted in panel **A**. The distance is measured between the geometrical centers of the quinone head group (atoms C_1_-C_6_), and the N2 iron-sulfur cluster. Three helices (TMH1, TMH3, and the lateral helix) of the ND1 subunit structures at the entrance to the UQ chamber. The cavity is plotted with the CAVER tool (v. 3.0) [36], taking ^49kDa^Tyr and ^49kDa^His as the reference with the cutoff radius of 0.7 Å.

**Fig. 4.**
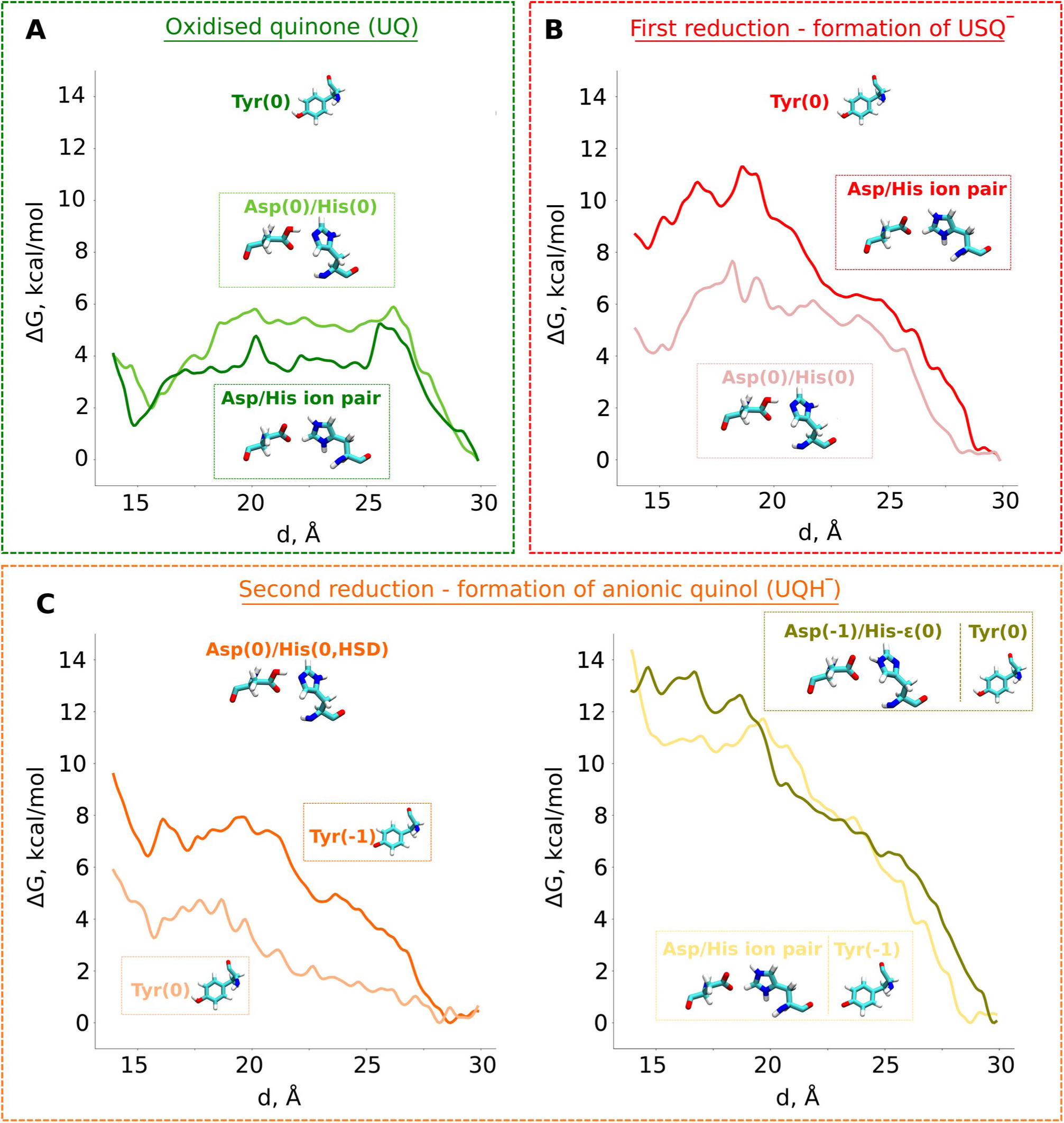
Classical free energy profiles of the propagation of UQ along the binding chamber of respiratory complex I. PMF profiles of (**A**) oxidised ubiquinone (UQ), (**B**) anionic semiubiquinone (USQ^-^), and (**C**) anionic ubiquinol (UQH^-^). The x-axis corresponds to the distance between the N2 iron-sulfur cluster and the quinone head group (Fig. 3B). The modelled protonation states of ^49kDa^Tyr (denoted as Tyr), ^49kDa^His (denoted as His), and ^49kDa^Asp (denoted as Asp) are listed in the legends.

Our QM/MM data revealed that protonic rearrangements in the active site lead to a stable state comprising charge-neutral histidine and aspartate residues (Fig. S2A). To understand the importance of different charge states of these residues on UQ binding, we performed classical free energy calculations in both protonation states of the His/Asp pair – by modelling both residues charged, and by modelling both residues charge-neutral. Contrary to the expected stable binding of anionic USQ with protonated histidine, we find His/Asp pair traps USQ^-^ better by ∼4 kcal/mol (Fig. 4B), when both residues are in their charge-neutral states. This unprecedented binding mode is in part due to the hydrogen bonds of anionic USQ with the titratable residues in the deep binding chamber (Fig. S3B). All in all, our data suggest that the anionic USQ radical remains trapped in the tunnel, with both histidine and aspartate maintained in their charge-neutral states.

### Two-electron reduction and formation of singly- and doubly-protonated ubiquinol

To study the two-electron reduction of UQ, we resorted to QM/MM simulations. Previous QM cluster models and short time scale QM/MM simulations showed that the two electron reduction results in local protonation of UQ by one or two protons [18,24]. Here, we performed free energy calculations to obtain quantitative estimates on the proton transfer dynamics from neighboring proton donors – the His/Asp system and tyrosine (Fig. 1, inset). We find that the ubiquinol anion (UQH^-^) formed after two-electron reduction is ca. 2 kcal/mol higher in energy than the doubly protonated UQH_2_ state, as long as the first proton transfer occurs from His/Asp pair (Fig. 2B, left panel). In contrast, proton transfer from the hydrogen bonded tyrosine creates a state with an ubiquinol anion that is much higher in energy (Fig. 2B, right panel, see also Fig. S4), presumably due to strong electrostatic repulsion between anionic tyrosine and UQH^-^. To further evaluate that QM/MM free energy calculation results (Fig. 2) do not depend on the size of the QM region, basis set, and the choice of the reaction coordinate, we performed additional calculations. Upon two-electron reduction of UQ with a larger QM region (Table S2), a similar free energy profile is obtained (Fig. S5B). Moreover, when changing the basis set from double-zeta (def2-SVP) to more accurate triple-zeta quality (def2-TZVP), the barrier separating UQH^-^ and UQH_2_ becomes higher, thereby kinetically stabilizing the partially protonated ubiquinol anion (Fig. S6).

Even though the energy difference between UQH_2_ and UQH^-^ is only ∼2 kcal/mol in favour of charge-neutral ubiquinol, this gives equilibrium occupancy of the neutral quinol 10-100:1. Therefore, to additionally probe the existence and stability of UQH^-^, we scrutinized our unbiased and biased classical and QM/MM MD simulation trajectories. Similar to the oxidized (UQ) and one-electron reduced (USQ^-^) state simulations, a shift in the position of histidine occurs also in the two-electron reduced state. It undergoes a conformational change from its structurally observed position to a direct hydrogen-bonding interaction with the ketone moiety of the UQ head group (Movie S2). However, in stark contrast to UQ and USQ^-^ states, a rapid proton transfer takes place from histidine to doubly-reduced UQ, concomitant with its reprotonation from neutral aspartate (Movies S1, S2) and leading to the formation of UQH_2_. Interestingly, this proton relay converts histidine from its one tautomeric state to another (from proton on δ nitrogen to proton on ε nitrogen). Analysis of long-timescale classical MD simulations showed a novel stable conformational arrangement of hydrogen bonding between histidine residue and methoxy groups of UQ (ca. 25% of simulation time, Fig. S3). Next, we initiated additional QM/MM setups starting from the classical MD snapshot in which histidine is stabilized in hydrogen-bonding with methoxy groups of UQ (see Table S2). In QM/MM MD simulations of all charge states (oxidized, one- and two-electron reduced), histidine remains “trapped” in between the quinone methoxy groups with hydrogen bonds, thereby preventing the delivery of the proton to doubly-reduced UQ, and resulting in the formation of UQH^-^ with a single proton donated by tyrosine (Movie S3).

### Multi-scale simulations explain the site-directed mutagenesis data

Assuming that the His/Asp pair and tyrosine are the only local proton donors to UQ upon its reduction, one would expect that their mutation to a non-polar residue would yield enzyme completely inactive. However, this is not the case [6]. Intriguingly, upon mutation of aspartate to asparagine, UQ reduction is not completely suppressed, instead, the enzyme still pumps protons, albeit sluggishly [18]. Similarly, mutation of tyrosine to phenylalanine renders the enzyme catalytically active with proton pumping [19,25]. Point mutation of histidine to arginine [25] also yields an enzyme that is still partially active. These data clearly show that single point mutation of putative proton donors to UQ is well tolerated and that upon two-electron reduction of UQ, proton must be transferred from the available donor (either tyrosine or Asp/His pair). Not surprisingly in our mutant QM/MM MD simulations, we observed that reduction-coupled proton transfer occurs either from Asp/His system or tyrosine leading to the formation of anionic ubiquinol (Fig. S2B,C). We next investigated the influence of point mutations by calculating the electron affinities of WT and mutant systems (see methods). Notably, the data suggests that first electron transfer is energetically comparable for both the WT and mutants (Fig. S7), however, for the second electron transfer (which is always coupled with proton transfer), the electron affinities of mutants are in part lower, commensurate with their lower catalytic activity and sluggish proton pumping [6]. These data suggest that UQH^-^ is most likely the species that forms during two-electron reduction and drives the catalysis of mutated complex I (see Discussion).

### Tunnel dynamics of anionic ubiquinol

To study the long-time scale behaviour of UQH^-^ species, we performed classical MD simulations in two different states representing proton donation either from tyrosine or from His/Asp pair (Fig. 3A). The high energy state representing anionic tyrosine (Fig. 2B, left panel) showed occupancy of anionic quinol at UQ binding sites 1 and 4 (Fig. 3A, brown curve). However, no such diffusion of UQH^-^ is observed in the limited time scales of unbiased MD simulations when proton transfer occurs from His/Asp system instead of tyrosine. In order to trigger the departure of UQH^-^ from site 1 and probe its energetics, we performed classical US MD simulations on several different states (Table S4). We find that UQH^-^ formed after single proton transfer is almost equally strongly repelled either by anionic tyrosine or by the negatively charged His/Asp pair (Fig. 4D, light yellow or olive-green traces, respectively). The obtained PMF profiles display almost barrierless UQH^-^ exit from site 1 towards site 4, in stark contrast to UQ and USQ^-^ cases representing oxidized and one-electron reduced UQ, respectively, where diffusion is impeded by kinetic barriers. In agreement with the data shown in Figs. 2 and S2A, the PMF profile in states with anionic tyrosine show 5-6 kcal/mol stabilization of neutral charge states of histidine and aspartate than their charged forms (Figs. 4, see also Fig. S8). As alluded to above, electrostatic repulsion between UQH^-^ and anionic tyrosine is likely the reason for the high energy of the state (Fig. 2), we simulated the exit of UQH^-^ from site 1 by modelling the neutral charge state of the tyrosine residue (Fig. 4C). The lowering of energy of UQH^-^ binding near N2 center by ca. 3 kcal/mol points to the key role of electrostatic interaction in driving anionic ubiquinol away from the site 1. A much stronger energetic stabilization is observed upon neutralization of either histidine or aspartate of His/Asp pair (net charge change from -1 to 0) with tyrosine modelled neutral (Fig. S8) further highlighting the importance of electrostatics in driving the UQH^-^ away from site 1 and towards site 4.

## Discussion

Based on our classical and QM/MM computations, we propose a stepwise mechanism of UQ reduction and its dynamics in the UQ tunnel. The proposed mechanism is in agreement with our recently developed model on the coupling mechanism of respiratory complex I [6], which we schematically illustrate in Fig. 5. The oxidised UQ binds at site 1 within the electron transfer distance to N2 FeS cluster. A kinetic barrier prevents the diffusion of UQ to site 4 (Fig. 5, top left panel). At this stage, both tyrosine and His/Asp pair are charge-neutral, with histidine and aspartate each holding one proton. Upon the first electron transfer to UQ, the semiquinone radical species is formed. The protonation of anionic semiquinone either from tyrosine or from His/Asp pair is unfavourable and it remains anionic in nature, which could be experimentally verified by EPR approaches. The binding of USQ^-^ at site 1 is less favorable compared to oxidized UQ, but the kinetic barrier towards site 4 prevents its escape (Fig. 5, top right panel) thereby enhancing the probability of its reduction by the second electron. The second electron transfer occurs concomitantly with a proton transfer from the His/Asp pair, where a proton relay converts histidine from its one tautomeric state to another (Fig. 5, bottom right panel), a notion that may be investigated with NMR experiments. The anionic ubiquinol formed (see also [26]) is strongly repelled by the now anionic His/Asp pair. At this stage, either deprotonation of tyrosine occurs to form UQH_2_ or UQH^-^ departs. The nearly isoenergetic states (UQH^-^ and UQH_2_) suggest reaction may proceed by involving either of the two species. In the case of the former, the free energy barrier is completely eliminated and there is a strong driving force for the diffusion of UQH^-^ towards site 4 (Fig. 5, bottom left panel). After reaching site 4, the anionic quinol drives the proton pump of respiratory complex I by proton injection-driven proton pumping mechanism [6]. Our results posit that both ubisemiquinone radical and ubiquinol are anionic in nature and their occupancy is likely enhanced in the mutants of the active site residues (histidine, tyrosine and aspartate). Thus, it would be desirable to look for spectroscopic signals in the mutated enzyme to strengthen the functional importance of anionic ubisemiquinone and anionic ubiquinol.

**Fig. 5.**
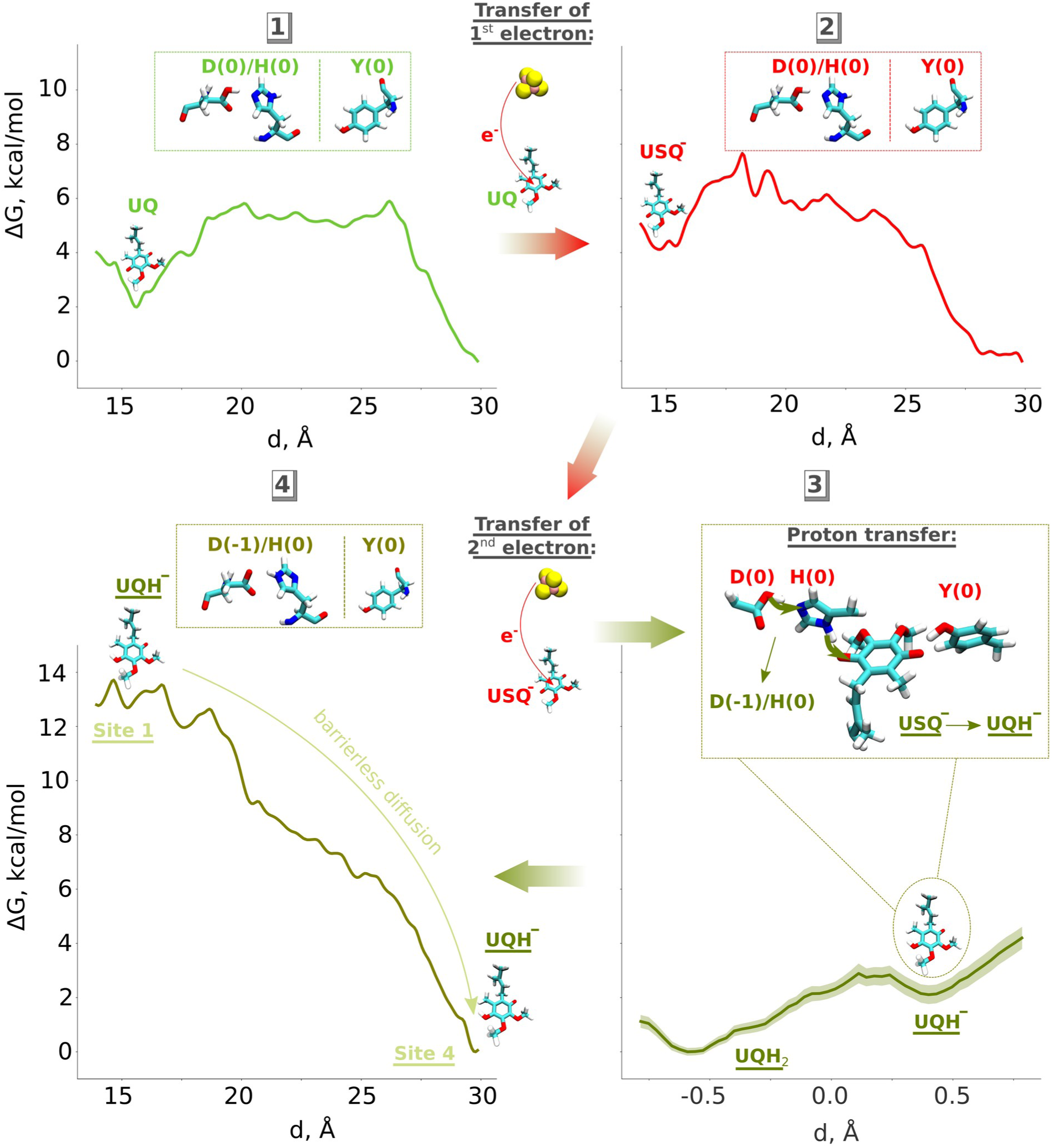
Proposed mechanism of ubiquinone reduction in respiratory complex I. Oxidised UQ (top left) gets an electron from the N2 iron-sulfur cluster to become anionic ubisemiquinone (USQ^-^, top right). Upon accepting the second electron from N2 (bottom left), a proton is transferred from the His/Asp pair leading to the formation of anionic ubiquinol (UQH^-^). The electrostatic repulsion by anionic His/Asp drives the diffusion of UQH^-^ towards site 4 of the UQ binding chamber.

### Computational methods Classical MD simulations

To study the long-timescale UQ dynamics in the UQ tunnel, we performed classical atomistic MD simulations of the high-resolution (2.4 Å) cryo-EM structure of respiratory complex I from *Yarrowia lipolytica* [27]. The model system comprised 8 core subunits: ND3, ND1, 49 kDa (excluding long terminus with residue IDs from 29 to 79), 30 kDa, PSST, TYKY, ND4L, and ND6, as well as 10 accessory subunits: 39 kDa, B13, B14.5a, B17.2 (excluding residue IDs 124-137), B14, PGIV, B16.6, 15 kDa, B9, and MWFE (Fig. S9). The protein system was embedded into a lipid bilayer consisting of POPC, POPE and TLCL lipids, with a periodic water/ion box of 145 Å x 160 Å x 200 Å, and NaCl salt concentration of 150 mM.

As no quinone molecule was resolved in the cryo-EM structure, we modelled UQ (nine isoprene units) species in the deep binding chamber by aligning the UQ coordinates taken from the complex I structure (PDBID 7V2C) [16]. We used different charge states of the UQ ligand: neutral (oxidised) state (UQ) [28], anionic ubisemiquinone (USQ^-^, charge -1), and anionic ubiquinol (UQH^-^, charge - 1). Parameters for UQ, USQ^-^, and UQH^-^ were taken from refs. [9,28,29]. With USQ^-^, we modelled the first step of quinone reduction, and with UQH^-^ the reduction by two electrons, and local protonation (either from ^49kDa^Tyr144 or from ^49kDa^His95/^49kDa^Asp196 pair). The system was first minimised for 1000 steps using the conjugate-gradient algorithm, as implemented in NAMD (v. 2.14) [30]. During the minimisation procedure, protein heavy atoms, quinone ring, and the lipid phosphorous atoms were harmonically restrained to their structural positions with the potential U=k(x-x_0_)^2^ and with a force constant of 99.0 kcal/(mol·Å^2^). After that, we performed minimisation in GROMACS (v. 2021.2) [31] using the steepest-descent algorithm with the force tolerance 5.97 kcal/(mol·Å^2^) (potential U=k/2 (x-x_0_)^2^), followed by a 10-ns equilibration in the NPT ensemble with the same constraints. Simulation replicas 2 and 3 were generated by extending the above equilibration procedure to 11 ns and 12 ns, respectively, therefore yielding a different starting conformation. The equilibration stage was concluded by a 10-ns NPT simulation with harmonic restraints on C_α_ atoms of the protein backbone. In the production run, all atoms were kept free. All classical MD simulations in this part were performed with GROMACS (v. 2021.2) software [31], using the V-rescale thermostat [32] and Berendsen barostat [33].

Classical MD umbrella sampling simulations were performed, starting from the last snapshot of the unbiased MD production run with UQ. In order to avoid convergence problems, we mutated UQ with 9 isoprenoid units to a UQ with a single isoprenoid unit, and then “pulled” the quinone from site 1 to site 4 with a velocity of 10^-7^ Å/fs, and a force constant of 10^3^ kJ/(mol·Å^2^). The reaction coordinate was represented by the distance between the geometrical centers of N2 iron-sulfur cluster and the quinone head group ring (Fig. 3B). Different umbrella sampling windows were generated by extracting the snapshots from the “pulling” run with a 0.5-Å stride between the reference values of the reaction coordinate. In order to alter the actual Q-N2 distance to the reference distance, each window was equilibrated for 10 ns with a larger force constant of the biasing potential (10^4^ kJ/(mol·Å^2^)). Then, we carried out a 40-ns equilibration with the lower force constant (10^3^ kJ/(mol·Å^2^)). The production simulation used for plotting free energy profiles was performed for 60 ns with a force constant of 10^3^ kJ/(mol·Å^2^). Histogram unbiasing procedure was implemented by WHAM analysis tool (v. 2.0.10) [34]. Setups for quinone in different charge states were created from the initial simulation snapshots of UQ MD simulation setup by converting quinone to the respective charge state in each of the simulation windows. The convergence plots of the PMF profiles for different time series are shown in Fig. S13.

Figures were prepared with VMD (v. 1.9.4) [35], and cavities were plotted with the CAVER (v. 3.0) software tool for protein analysis and visualisation [36].

### Hybrid QM/MM simulations

Hybrid quantum-mechanical/molecular-mechanical (QM/MM) MD simulations were performed on the high-resolution structure of mitochondrial complex I from *Sus scrofa* [16], where the UQ ligand has been fully resolved in the UQ tunnel (Fig. S10, inset). Six core subunits were included in the model system: ND3, ND1, 49 kDa, 30 kDa, PSST, and TYKY. To eliminate the large spatial fluctuations of the long N-terminus of the 49kDa subunit, residues 34 to 78 were excluded from the model system.

To study the protonation dynamics of the oxidized and reduced UQ species, we first carried out unbiased QM/MM MD simulations of quinone ligand structurally resolved at site 1 with a direct hydrogen bond with the highly conserved Tyr141 of the 49 kDa subunit (Fig. 1, inset). The QM region comprised protein sidechains surrounding the UQ head group (Table S1). For QM/MM boundary, protein amino acids were pruned at the C_α_-C_β_ bond, while the QM/MM bond for Q_9_ was created between the C_11_ and C_12_ atoms. Linking hydrogens were placed along the QM/MM bonds, as implemented in NAMD QM/MM module [37]. Hydrogen atoms were added to the structural conformation using the PSFGEN plugin for VMD [35], and the protonation states of the titratable residues were derived from the pKa calculations [38] performed by PROPKA software [39]. The system was embedded into a lipid bilayer containing 50% POPC, 36% POPE, and 14% cardiolipin (TLCL) ligands, and immersed into a water-ion box with the dimensions 110x124x125 Å^3^, and with the 150 mM concentration of NaCl salt. In total, the simulation system comprised around 211,000 atoms.

The simulations were carried out with NAMD molecular dynamics package (v. 2.14) [30,37] in combination with ORCA quantum chemistry program (v. 5.0.3) [40]. The MM part was treated with the CHARMM36 force field [41], where the non-bonded interactions cutoff was set to 12 Å, with switching and pairlist distances equal to 10 Å and 14 Å, respectively. The system was classically minimised for 1000 steps with the conjugate-gradient algorithm, followed by a 10-ns classical MD equilibration in the NPT ensemble with harmonic restraints on all heavy atoms of protein and quinone (with a force constant of 99.0 kcal/(mol·Å)). In order to “relax” the system on its free energy landscape, an additional classical 10-ns equilibration stage with constraints on protein backbone was performed in certain setups (Table S2). This was followed by the QM/MM minimisation for 200 steps, and the unbiased QM/MM MD run. Temperature was maintained at 310 K using Langevin thermostat [42], and the pressure control was implemented with the Nosé-Hoover Langevin piston barostat [43,44]. QM region was described by density functional theory (DFT) with hybrid B3LYP functional [45] and def2-SVP basis set [46]. The DFT-D3 dispersion correction [47] was used to account for the long-range London dispersion forces, and the energy tolerance for the self-consistent field procedure was set to 10^-8^ au. The additive electrostatic embedding scheme [37] was used to impart Coulomb interactions between the QM and MM regions.

To probe the energetics of proton transfer, we carried out free energy umbrella sampling simulations. To facilitate the proton transfer reaction, the reaction coordinate was chosen as a linear combination of the distances forming the proton pathway (Fig S11):

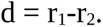

The first simulation window was created by applying a harmonic potential to this reaction coordinate, restraining it to its equilibrium distance:

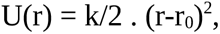

with the force constant k = 100 kcal/(mo·Å^2^). After that, the equilibrium distance r_0_ was moved by 0.02 Å every step to create subsequent windows with the stride of 0.2 Å. The bias to the reaction coordinate was introduced with the flexible *Colvars* module [48], implemented in NAMD. Each window was simulated for 3.8 ps in total, where the last 2 ps were taken for the calculations of the potential of mean force (PMF) profiles. These calculations were done using the weighted histogram analysis method with the WHAM (v. 2.0.10) analysis tool [34]. The bootstrapping errors were calculated using the correlation time of 20 fs and are depicted as a shaded area on the free energy profiles (Figs. 2, S4-S6, S14).

To further test the choice of the reaction coordinate, we performed additional US simulations by including the linear combination of His-Asp distances to the reaction coordinate for the UQH _2_-His proton transfer in a two-electron reduced state (Fig. S14, top inset). We show that the different reaction coordinate does not induce the formation of a metastable UQH^-^/His-ε(0)/Asp(0)/Tyr(-1) state (Fig. S14), coherent with our result for the UQH_2_-His reaction coordinate (Fig. 2).

Electron affinity for the first electron (Fig. S7) was calculated as an energy difference between the oxidised ubiquinone (E_UQ_) and ubisemiquinone (E_USQ_):

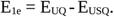

Similarly, the electron affinity for the second electron was derived as an energy difference between ubisemiquinone (E_USQ_) and ubiquinol (E_UQH_):

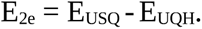

Energies were calculated as averages between 501^st^ and 1500^th^ fs of unbiased MD. Standard deviations for electron affinities σ_1e_, σ_2e_ were calculated as the square-root of the sum of squares of deviations of the respective individual distributions:

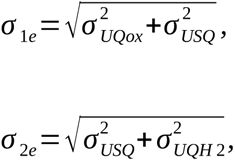

where σ_UQox_, σ_USQ_, σ_UQH2_ are the standard deviations on the QM/MM energy distributions in the zero-, one-, and two-electron reduced states, respectively.

## Authors contributions

OZ performed QM/MM MD simulations, classical MD simulations, free energy calculations, analysed data, prepared figures, co-supervised JL, and wrote the initial draft of the manuscript. JL performed QM/MM MD simulations. VS designed and supervised the research, analysed data, wrote and refined the text.

## Acknowledgements

VS acknowledges research funding from the Research Council of Finland, the Jane and Aatos Erkko Foundation, the Sigrid Jusélius Foundation, the Cancer Foundation and the University of Helsinki. VS and OZ thank the Center for Scientific Computing (CSC), Finland for providing large-scale computational resources. The Faculty of Science, University of Helsinki computational support is acknowledged. VS and OZ are thankful to Dr. Jonathan Lasham and Amina Djurabekova for fruitful discussions, help with classical MD simulations, and proofreading the manuscript. VS and OZ acknowledge helpful discussions with Dr. Outi Haapanen, Prof. Volker Zickermann, Dr. Giray Enkavi and Prof. Thorsten Friedrich.

