## Supplementary Information for "Catalytic relevance of quinol anion in biological energy conversion by respiratory complex I"

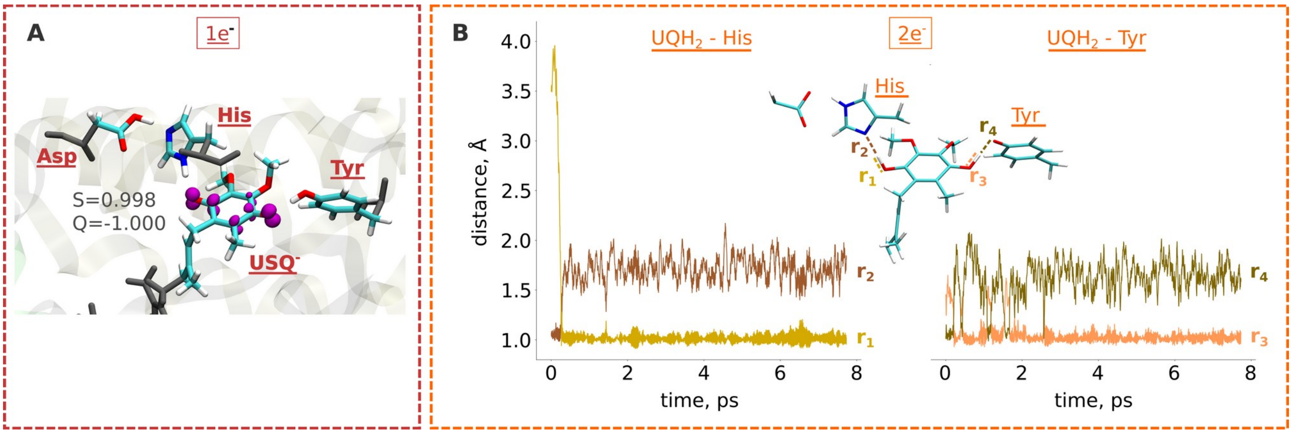


Fig. S1. **A** Spin density distribution (shown as magenta isosurface, isovalue 0.01) from the USQ model system obtained from the 1000$^{th}$ fs of the unbiased QM/MM MD simulation. Captions indicate total charge (Q) and spin (S), calculated with Mulliken population analysis. **B** Protonation dynamics in the two-electron reduced state observed in our unbiased QM/MM MD simulations. Hydrogen bond distances between the quinone ketone group and His (left panel) and Tyr (right panel). The distance notations are introduced in the inset.

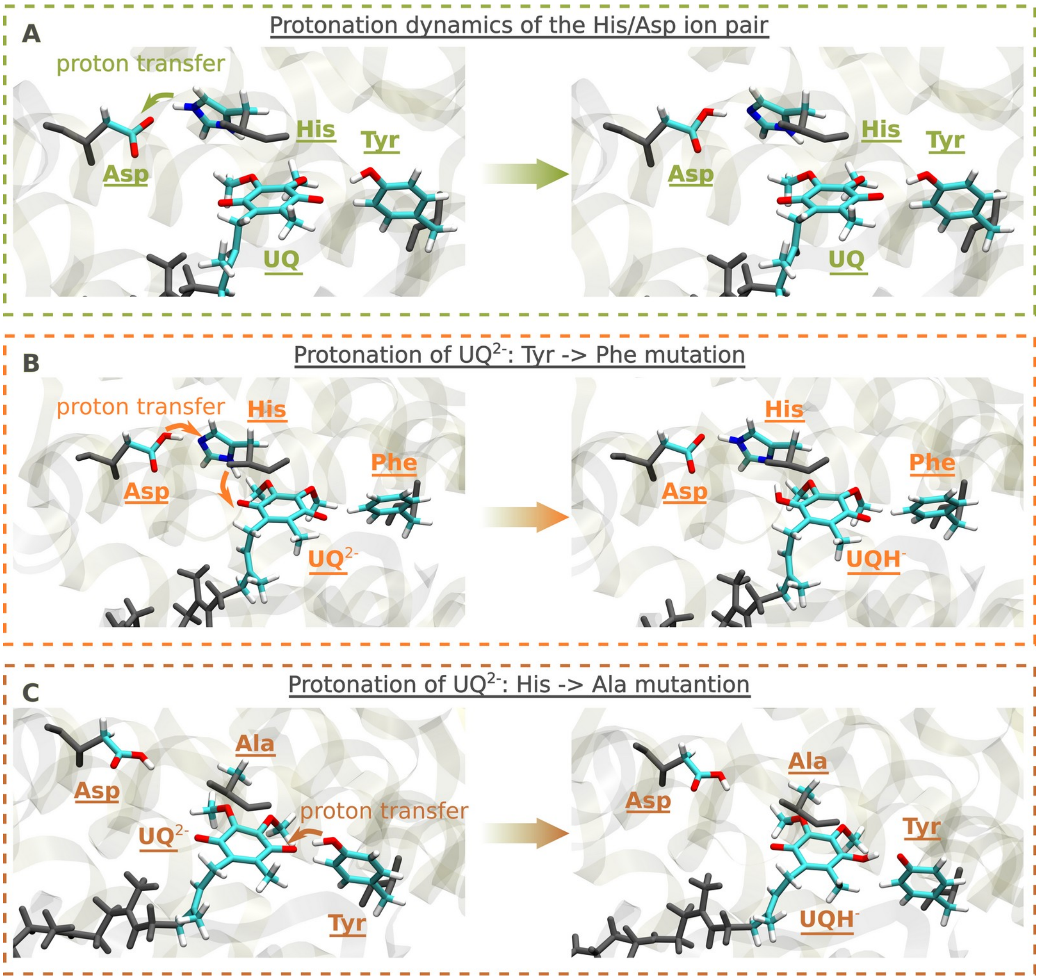


Fig. S2. Spontaneous protonation dynamics in unbiased QM/MM simulations: (**A**) Internal proton transfer within the His/Asp pair during the QM/MM minimization in oxidized and one-electron reduced states. (**B**) Proton transfer from the His/Asp pair to UQ during unbiased QM/MM MD simulations of two-electron reduced system, where Tyr is mutated to Phe. (**C**) Protonation of doubly-reduced ubiquinone from Tyr during unbiased QM/MM MD simulation, where His is mutated to Ala.

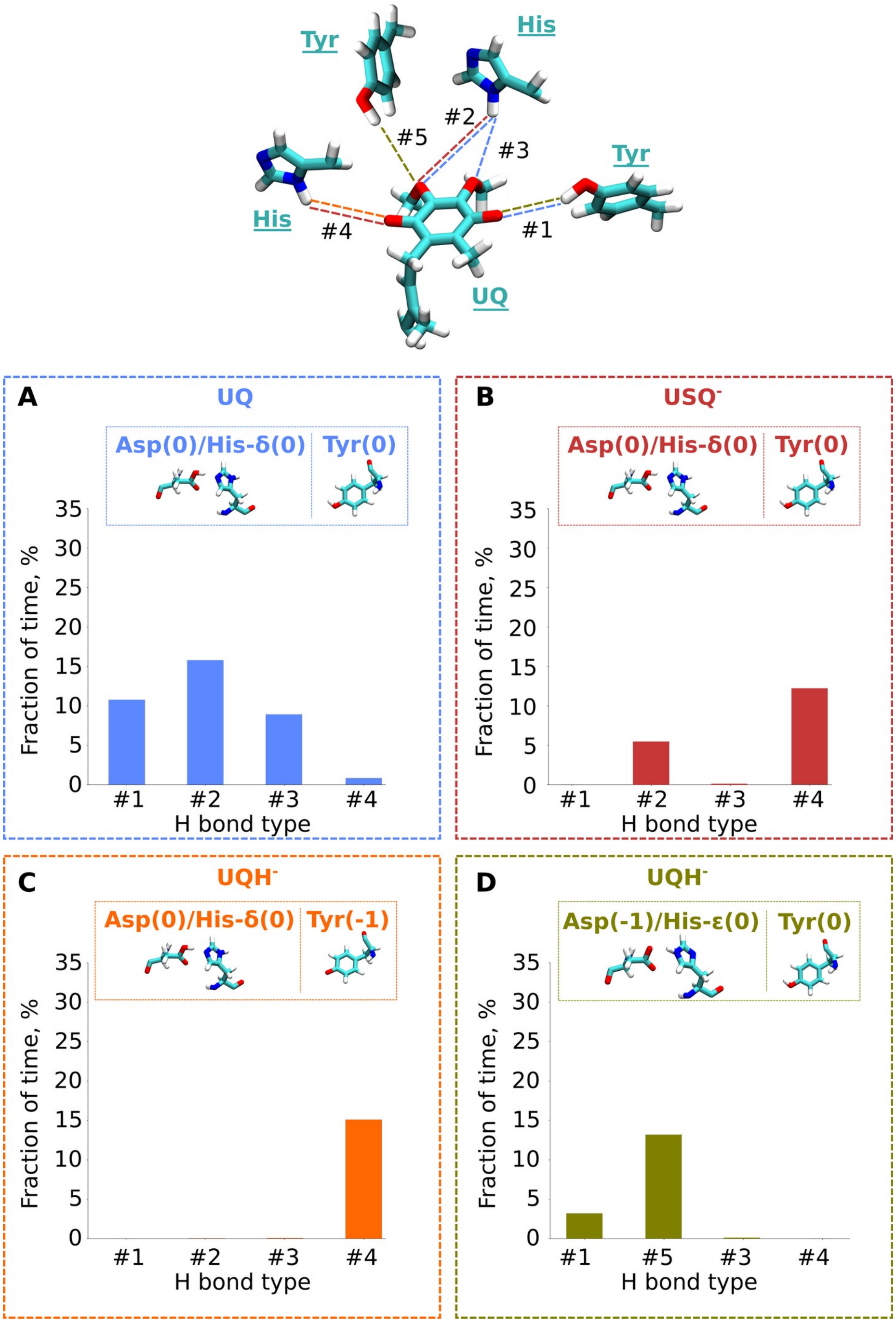


Fig. S3. Formation of hydrogen bonds by the UQ head group with the conserved residues at UQ binding site 1, obtained from classical unbiased MD simulations. Upper panel illustrates the types of hydrogen bonds listed on the x-axis of panels (**A**) – (**D**). Colors of the dashed lines denote the hydrogen bonds which are prevalent (exist more than 2.5% of the simulated time) in the respective setups (**A**) – (**D**). Insets on panels (**A**) - (**D**) indicate the modeled protonation states of the titratable residues, as well as UQ charge state.

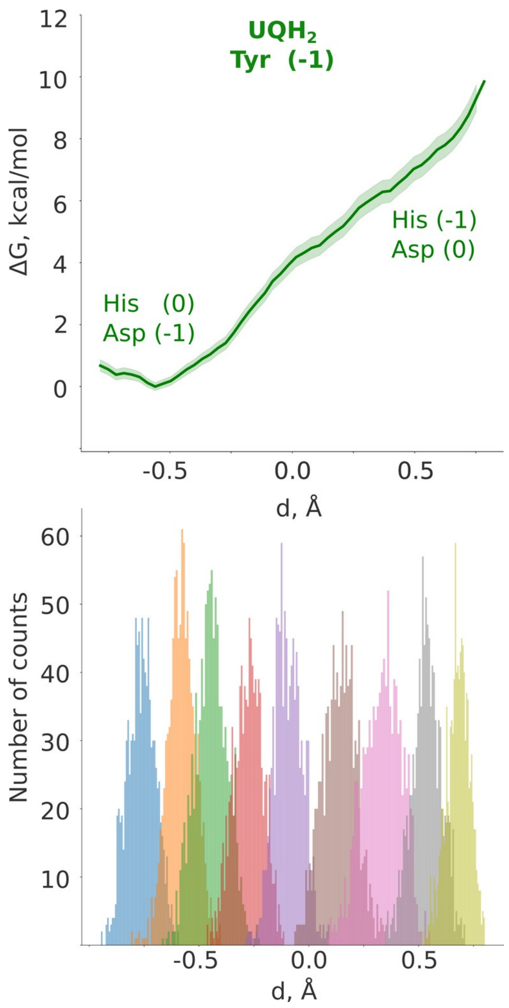


Fig. S4. QM/MM umbrella sampling free energy calculations of the proton transfer from His to Asp in the two-electron reduced case. Free energy profile (top panel), and occupational histograms of the reaction coordinate (bottom panel) sampled from the simulation windows.

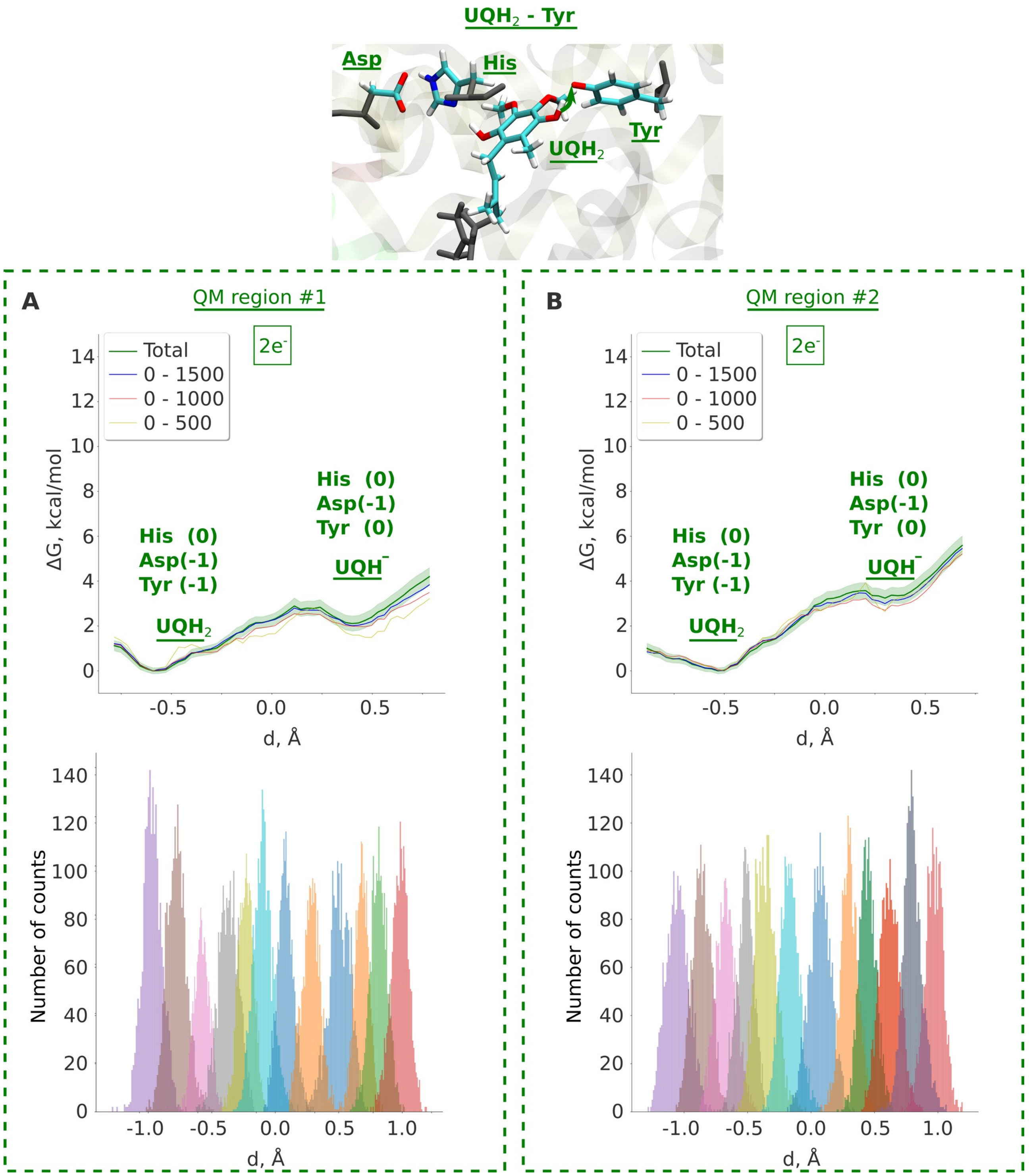


Fig. S5. Comparison between the results from QM/MM umbrella sampling simulations of the proton transfer from $UQH_2$ to Tyr in different QM regions (see Table S1): (**A**) QM region #1, results are also shown in Fig. 2B (right curve) of the main text. (**B**) Larger QM region (QM region #2). The captions to each energy state indicate the charge of the respective surrounding residues. Top panels show free energy profiles, and the bottom ones – occupational histograms of the reaction coordinate in the respective simulation windows.

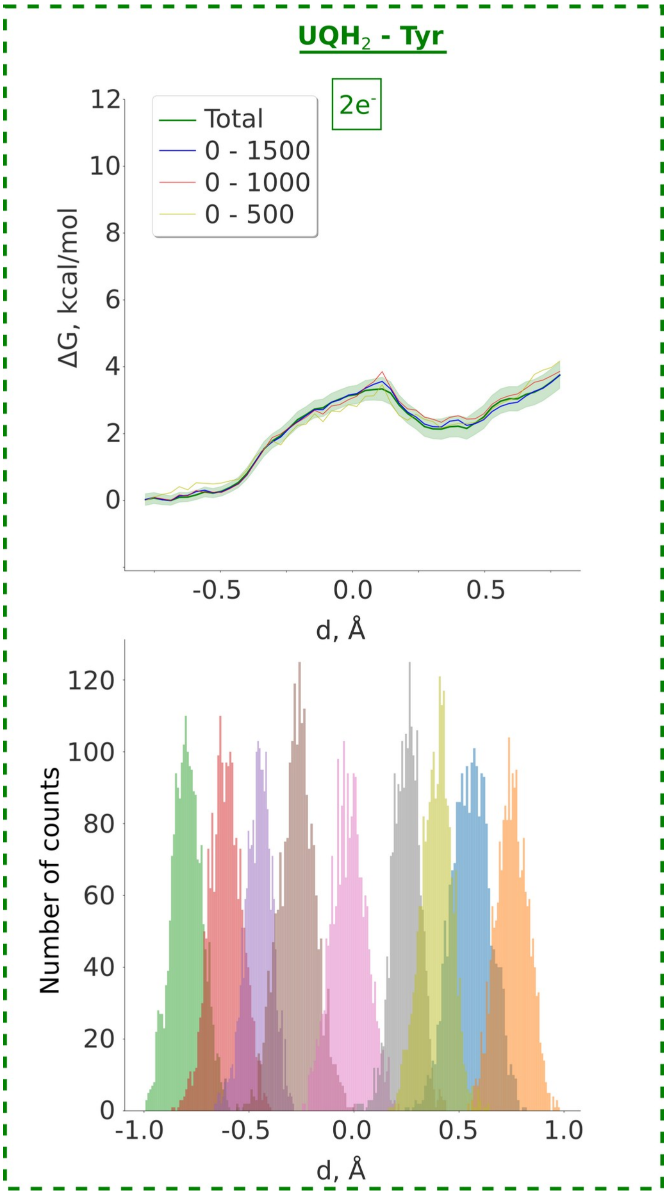


Fig. S6. Free energy profile (top) and occupational histograms (bottom) of the proton transfer from $UQH_2$ to Tyr using split-valence triple-zeta def2-TZVP basis set. Partial contributions to the free energy profiles from the first 500 fs, 1000 fs, and 1500 fs are shown as brown, red, and blue lines, respectively, highlighting convergence.

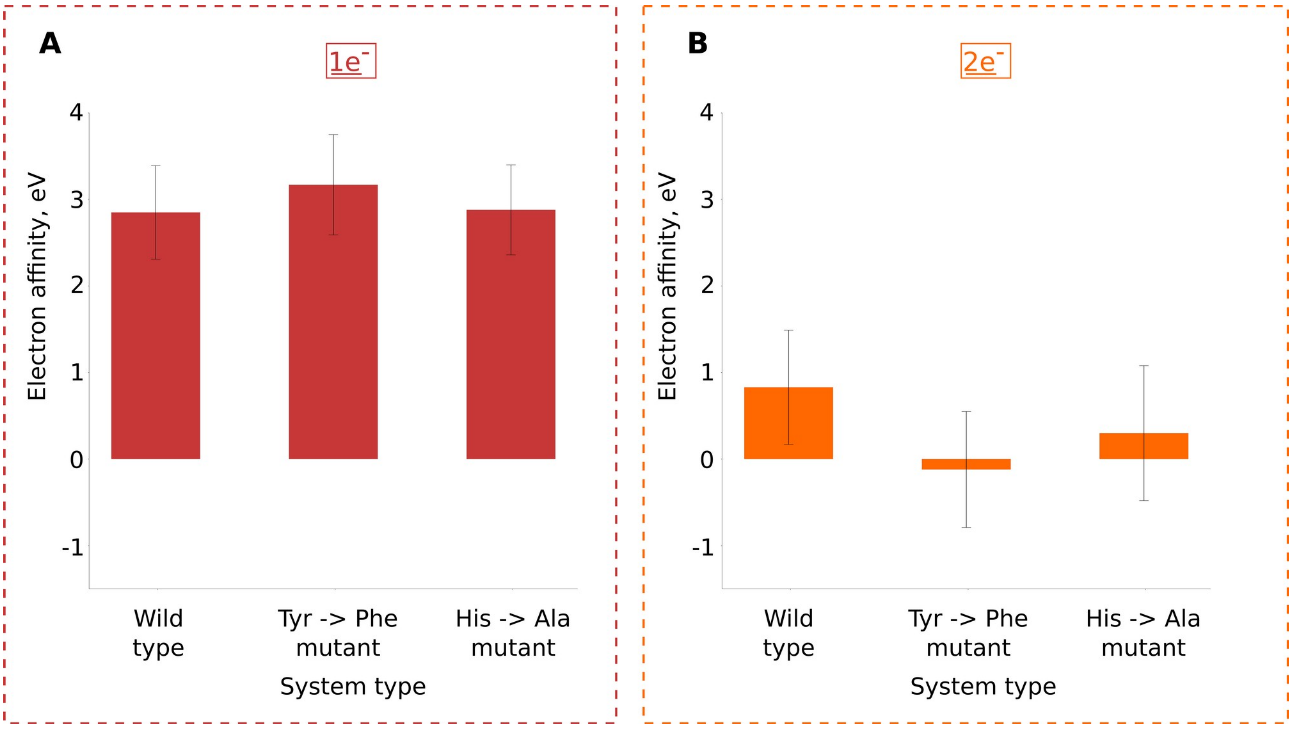


Fig. S7. Electron affinities for first (1e$^-$) and second (2e$^-$) electron transfer, calculated based on the QM/MM energies (see methods). Proton-coupled electron transfer upon adding one and two electrons to the system is illustrated in Fig. S2 B,C.

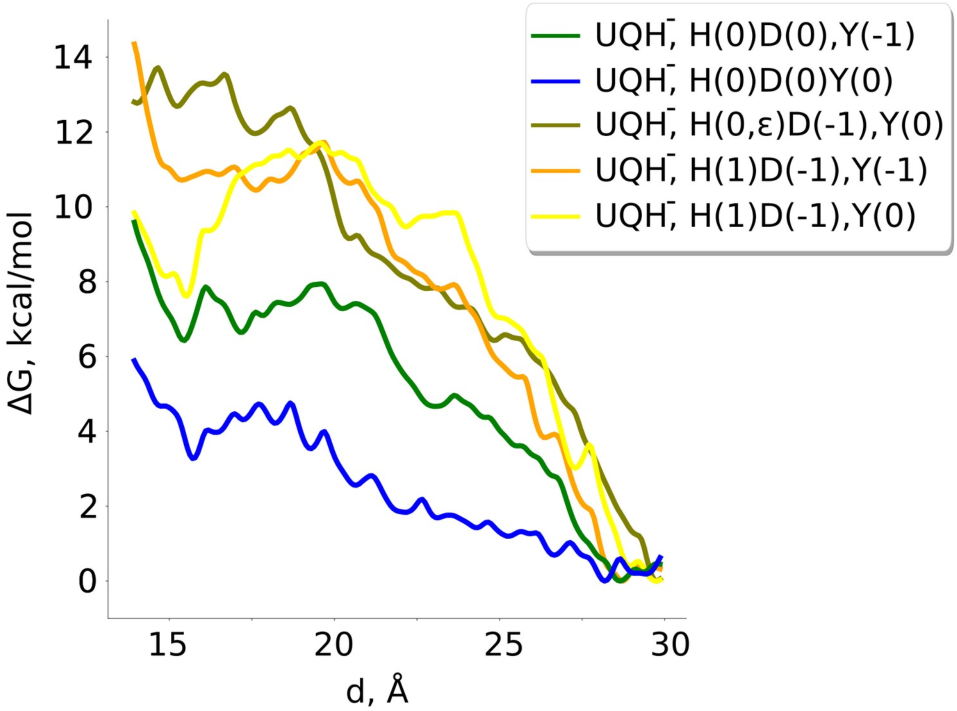


Fig. S8. Classical free energy profiles of the anionic ubiquinol species in the binding chamber of respiratory complex I with different charge states of the adjacent titratable residues: neutral His/Asp pair with anionic Tyr (green line); neutral His/Asp pair with neutral Tyr (blue line); neutral His with negatively charged Asp and neutral Tyr (olive-green line); His/Asp ion pair with anionic Tyr (orange line); His/Asp ion pair with neutral Tyr (yellow line).

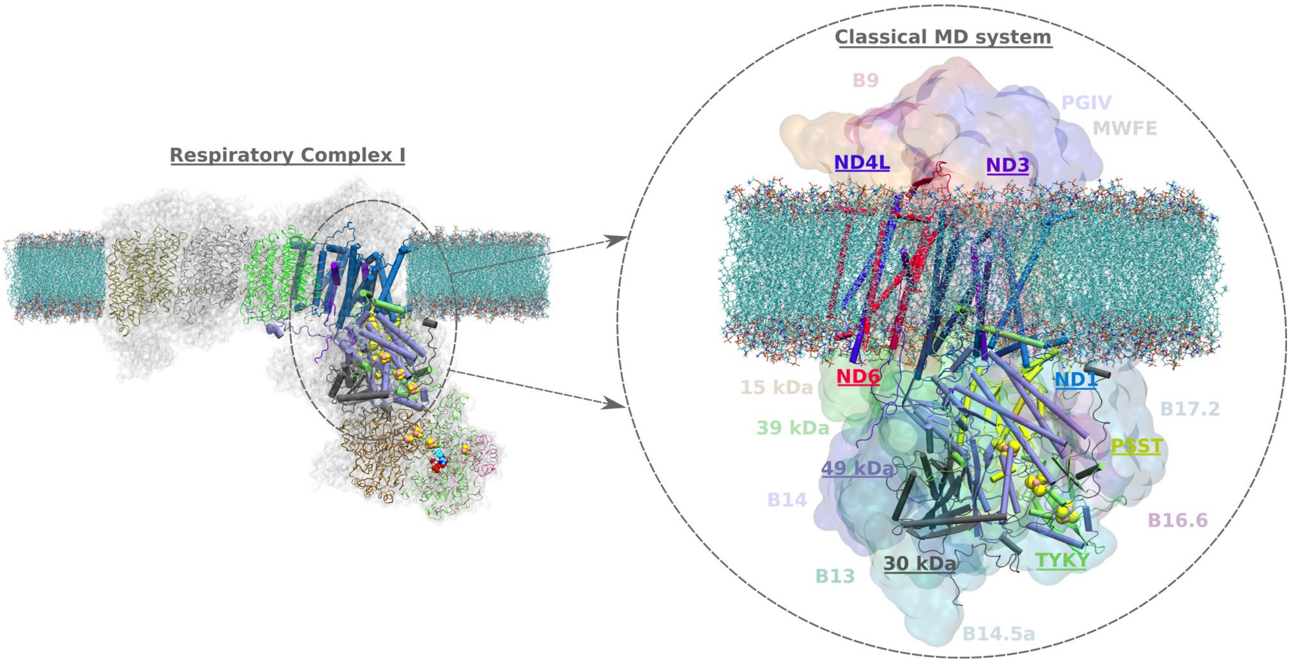


Fig. S9. Model system for classical MD simulations (right panel) constructed from high-resolution structure of mitochondrial complex I from *Yarrowia lipolytica* (left panel, PDB ID: 7O71) [1]. Cartoon representation in the right inset depicts the core subunits included in the model system. The accessory subunits are shown in surface representation.

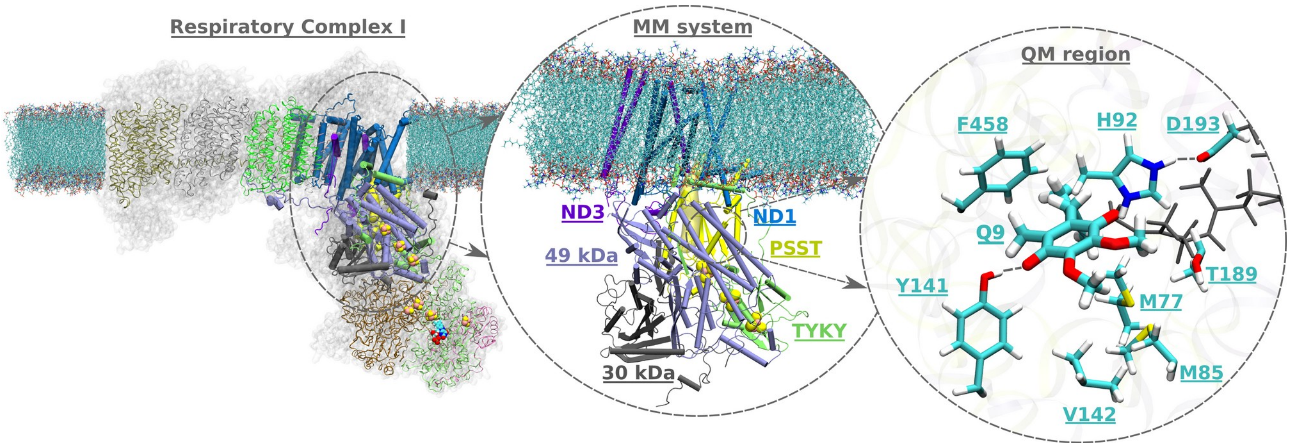


Fig. S10. Model system for hybrid QM/MM simulations of high-resolution structure of mitochondrial complex I from *Sus scrofa* (PDB ID: 7V2C [2]. Middle inset depicts the MM system consisting of 6 core subunits shown in respective colors: ND1, ND3, 49 kDa, 30 kDa, PSST, and TYKY. Water box and ions, which are also included in the MM region are not shown for clarity. Right inset illustrates protein residues and quinone head group included in the QM region. Hydrophobic isoprenyl tail of UQ included in the MM region is shown in grey licorice representation.

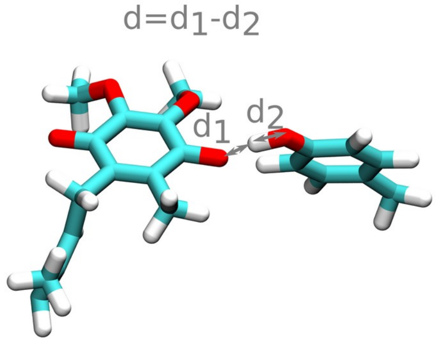


Fig. S11. An example reaction coordinate used in the QM/MM MD umbrella sampling simulations.

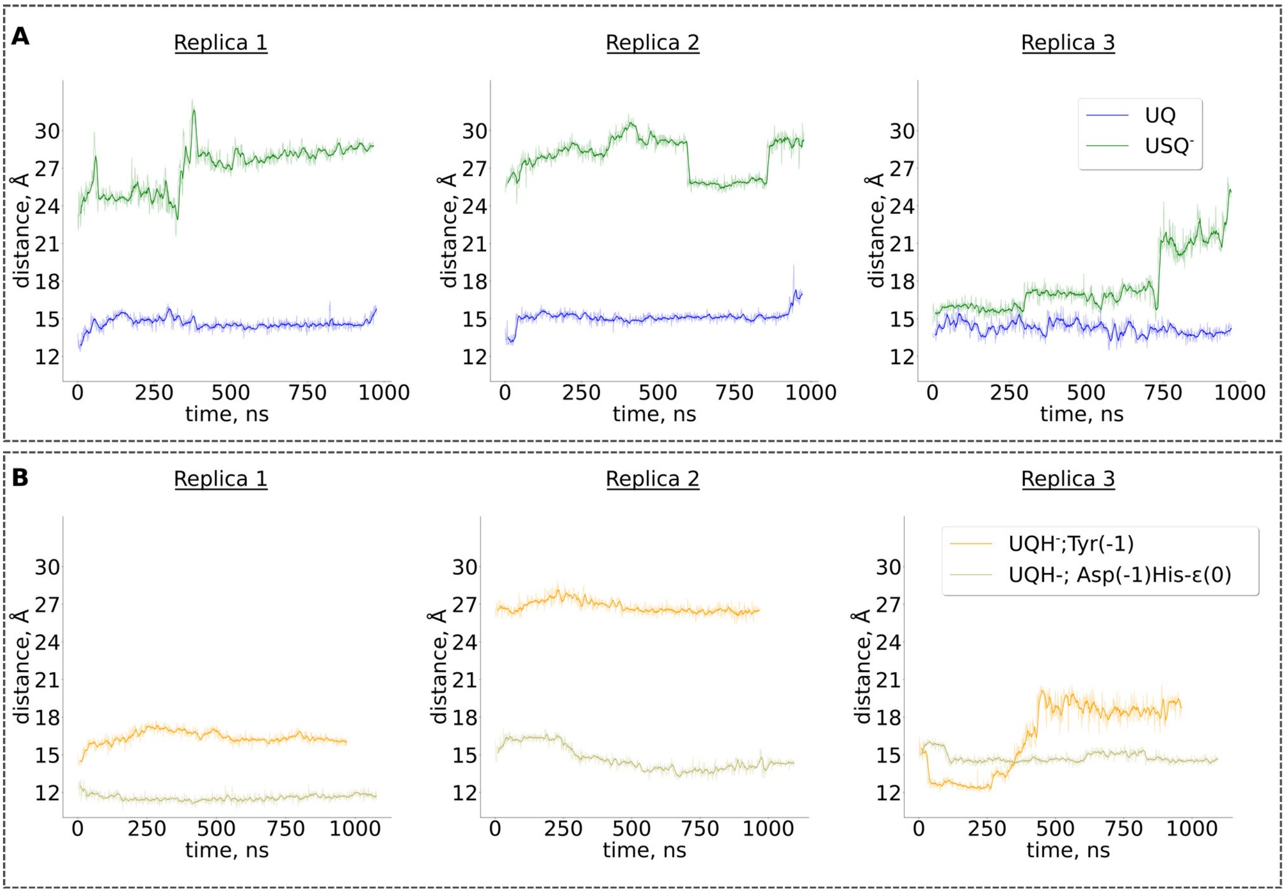


Fig. S12. Time series of the UQ-N2 distances (Fig. 3) from unbiased classical MD simulations for different charge states of the UQ species: **A** oxidised quinone (UQ, blue) and anionic ubisemiquinone (USQ$^-$, green). **B** anionic ubiquinol (UQH$^-$) with deprotonated Tyr (orange curve), and with deprotonated Asp and neutral His with the ε nitrogen protonated (brown curve).

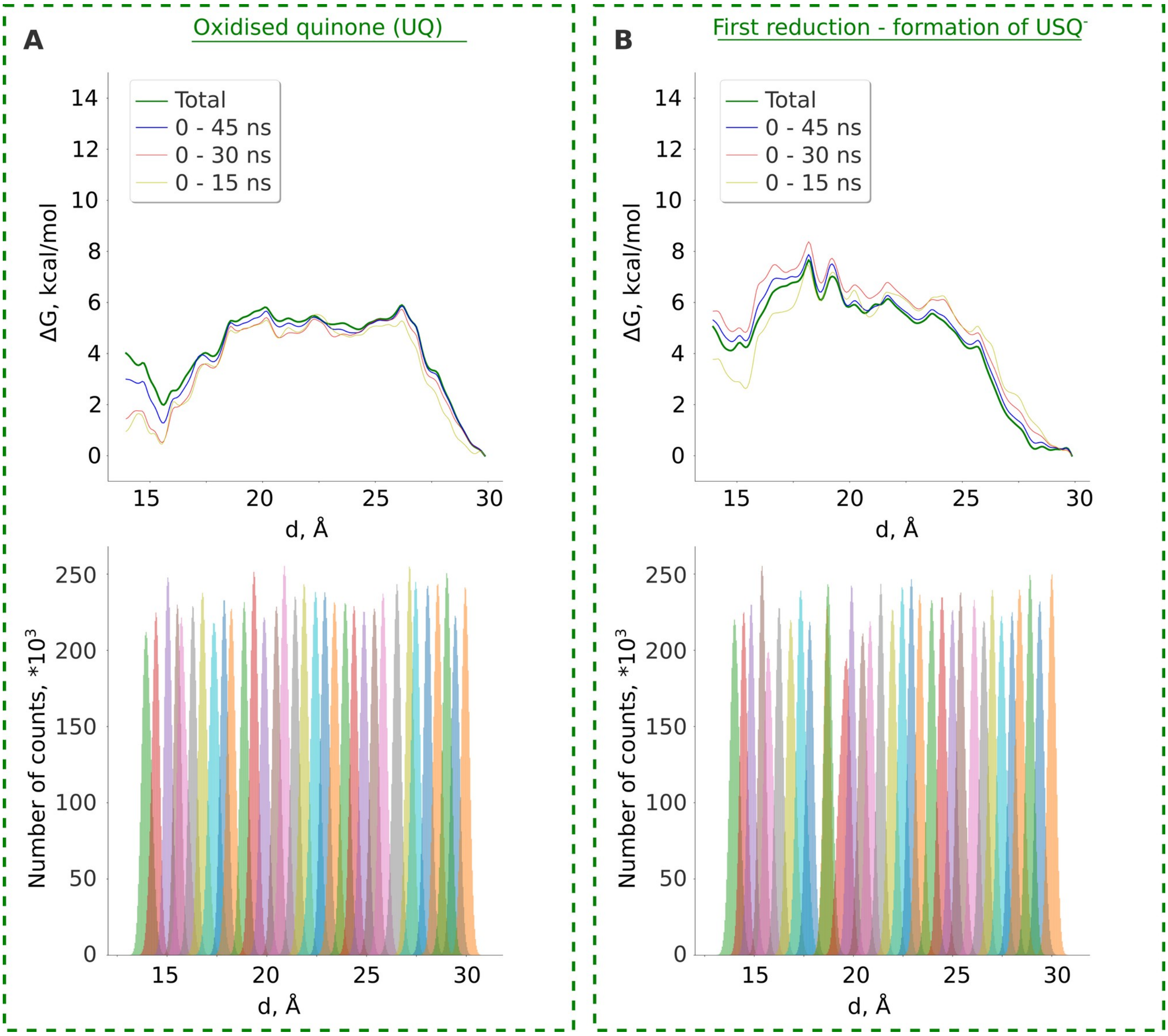


Fig. S13. Classical US free energy calculations of the oxidised UQ (**A**) and anionic USQ (**B**) in the quinone binding chamber. Top panels show free energy profiles, while the bottom panels depict the respective occupational histograms. Partial contributions to the total free energy profile from the first 15, 30, and 45 ns are shown with brown, red, and blue lines, respectively.

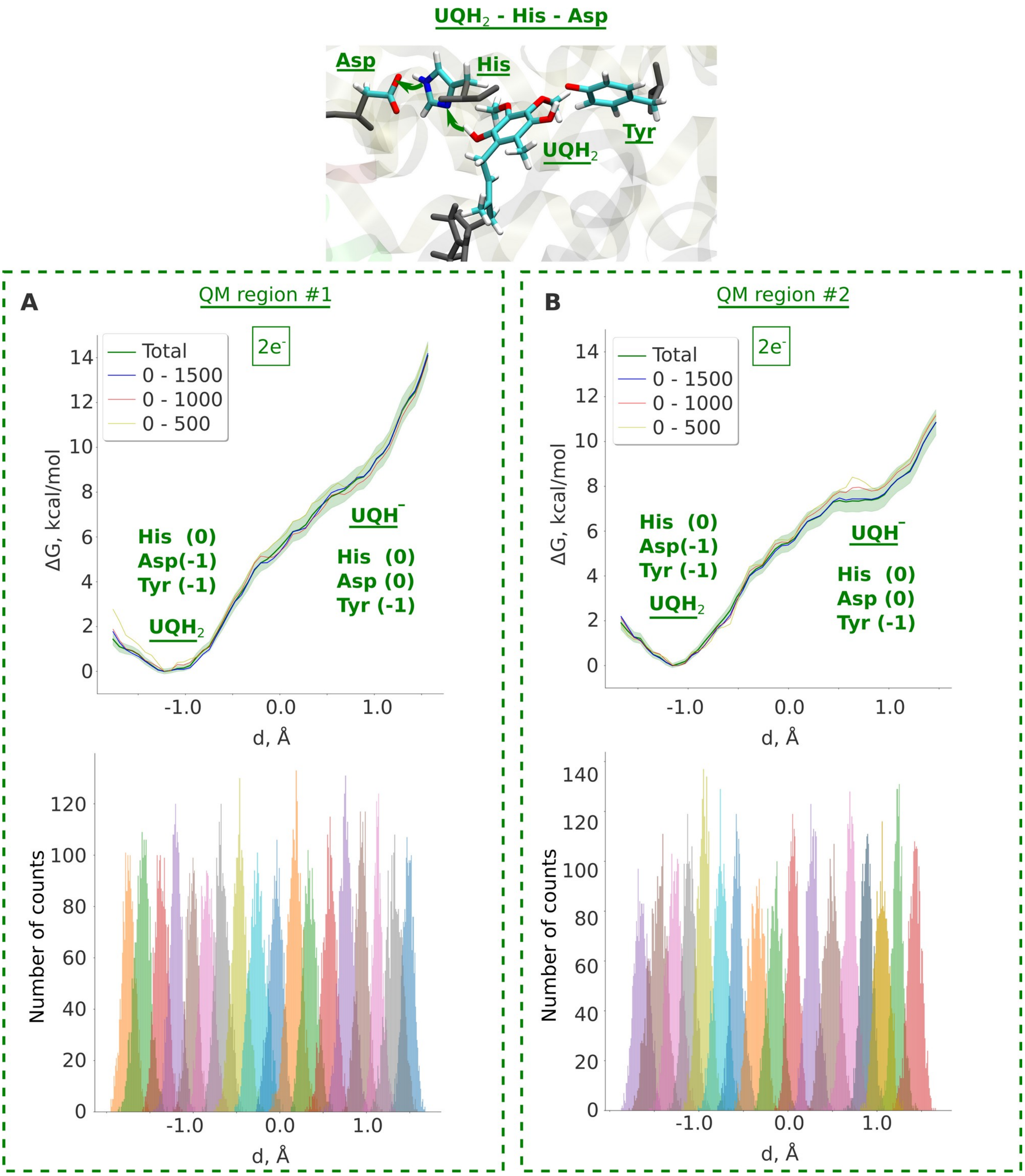


Fig. S14. Free energy profiles obtained from QM/MM umbrella sampling simulations of the proton transfer from $UQH_2$ to the His/Asp pair, where both hydrogen bonds ($UQH_2$..His and His...Asp) are included in the reaction coordinate (see inset). QM region #1 (**A**) and QM region #2 (**B**) (for QM region notations, see Table S1). Charges of the surrounding residues are depicted in captions to the respective energy states. Bottom panels show normalized distributions of the reaction coordinate in simulation windows.

Movie S1. Conformational rearrangement of His during the unbiased QM/MM MD simulation of a one-electron reduced state. The trajectory length included in the movie is 2.0 ps.

Movie S2. Protonation dynamics of ubiquinone during the unbiased QM/MM MD simulation in a two-electron reduced state. The movie is rendered from a 2.0-ps trajectory.

Movie S3. Formation and chemical stability of the quinol anion in the two-electron reduced system where quinone methoxy groups were initially symmetrically arranged with respect to each other. The video includes a 200-steps QM/MM minimization, as well as the full 3.7-ps dataset of unbiased QM/MM MD (see Table S2).

Table S1. QM regions considered in this work.

| QM region number | Starting structure (PDBId) | Number of atoms (excluding link hydrogens) | Residues included in the QM region |
|---|---|---|---|
| 1 | 7V2C [2] | 122 | $^{49kD}$Asp193, $^{49kD}$His92, $^{49kD}$Tyr141, $^{49kD}$Met185, $^{49kD}$Phe458, $^{49kD}$Thr189, $^{49kD}$Val142, $^{PSST}$M77, Q9 head group (QM/MM bond along C11-C12) |
| 1.1 | 7V2C | 131 | $^{49kD}$Asp193, $^{49kD}$His92, $^{49kD}$Tyr141, $^{49kD}$Met185, $^{49kD}$Phe458, $^{49kD}$Thr189, $^{49kD}$Val142, $^{PSST}$M77, Q9 head group (QM/MM bond along C11-C12), 3 water molecules aligned from the high-resolution structure from *Bos taurus* (PDB ID: 8Q48) [3] ($^{49kD}$Wat606, $^{49kD}$Wat643, $^{49kD}$Wat695) |
| 2 | 7V2C | 150 | $^{49kD}$Asp193, $^{49kD}$His92, $^{49kD}$Tyr141, $^{49kD}$Met185, $^{49kD}$Phe458, $^{49kD}$Thr189, $^{49kD}$Val142, **$^{49kD}$Lys404**, **$^{49kD}$Asn182**, **$^{49kD}$Ile456**, $^{PSST}$M77, Q9 head group (QM/MM bond along C11-C12) |
| 3 | 7O71 | 162 | $^{49kD}$Asp196, $^{49kD}$His95 $^{49kD}$Tyr144, $^{49kD}$Met188, $^{49kD}$Phe461, $^{49kD}$Thr189, $^{49kD}$Ser192, $^{49kD}$Phe203, $^{49kD}$Met195, $^{49kD}$Val145, $^{49kD}$Val97, $^{PSST}$Met91, $^{PSST}$Ala87, Q9 head group (QM/MM bond along C11-C12), 6 water molecules populating the QM region in the unbiased classical MD simulation‡ |

‡ the starting configuration for the QM/MM setups was taken from the classical MD snapshot after 300 ns of the production run (see Table S2)

Table S2. Wild-type QM/MM MD simulation setups constructed from the structures (PDBIds 7V2C and 7O71) [1,2].

| Initial structure (PDBId) | Equilibration protocol[1] | Extra electrons | QM/MM min | QM/MM MD | QM region (see Tabl. S1) | Umbrella sampling | Basis set |
|---|---|---|---|---|---|---|---|
| 7V2C | Std | 0e- | + | 2.0 ps | 1 | - | def2-SVP |
| 7V2C | Std | 1e- | + | 2.0 ps | 1 | UQ-Tyr141 | def2-SVP |
| 7V2C | Std | 2e- | + | 7.8 ps | 1 | UQ-Tyr141 | def2-SVP, def2-TZVP |
| | | | | | | UQ-His92<br>His92-Asp193<br>UQ-Tyr141 with constraints on His92(NE2-HE2) | def2-SVP |
| 7V2C | Std | 2e- | + | 7.8 ps | 2 | UQ-Tyr141 | def2-SVP |
| 7V2C[#] | Std | 0e- | + | 2.1 ps | 1.1 | - | def2-SVP |
| 7V2C[#] | Std | 2e- | + | 1.4 ps | 1.1 | - | def2-SVP |
| 7V2C[&] | Std | 2e- | + | 1.2 ps | 1.1 | UQ-Tyr | def2-SVP |
| 7V2C[*] | Std + Add | 0e- | + | 3.7 ps | 1 | - | def2-SVP |
| 7V2C[*] | Std + Add | 2e- | + | 3.7 ps | 1 | - | def2-SVP |
| 7O71[‡] | Std | 0e- | + | 1.2 ps | 3 | - | def2-SVP |
| 7O71[‡] | Std | 2e- | + | 1.64 ps | 3 | - | def2-SVP |

[1] “Std” stands for “standard” equilibration protocol, while “Std+Add” introduces an additional 10-ns equilibration stage with constraints on protein backbone, see the “Computational methods” section.

[*] one of two ubiquinone methoxy groups ($CH_3$-) was rotated to achieve symmetrical conformation of the UQ head group

[#] the setup was created by aligning 3 water molecules ($^{49kD}$Wat606, $^{49kD}$Wat643, $^{49kD}$Wat695) from [3] (PDB ID: 8Q48), located in the vicinity of the QM region, to the 7V2C structure

[&] the setup was created by aligning 3 water molecules ($^{49kD}$Wat606, $^{49kD}$Wat643, $^{49kD}$Wat695) from [3] (PDB ID: 8Q48), located in the vicinity of the QM region, to the 7V2C structure. The starting snapshot for QM/MM runs was taken after 1 ps of unbiased QM/MM MD of the "Std" 7V2C setup with 2 additional electrons

[‡] the starting configuration for the QM/MM setups was taken from the classical MD snapshot after 300 ns of the production run

Table S3. QM/MM MD simulation setups with introduced mutations (Tyr/Phe and His/Ala) constructed from the structure (PDBId: 7V2C).

| Initial structure (PDBId) | Equilibration protocol[1] | Extra electrons | QM/MM min | QM/MM MD | QM region (see Tabl. S1) | Umbrella sampling | Basis set |
|---|---|---|---|---|---|---|---|
| **Mutant: Tyr144 -> Phe** | | | | | | | |
| 7V2C | Std | 0 | + | 2.75 ps | 1[2] | - | def2-SVP |
| 7V2C | Std | 1e- | + | 1.97 ps | 1[2] | - | def2-SVP |
| 7V2C | Std | 2e- | + | 3.4 ps | 1[2] | - | def2-SVP |
| **Mutant: His92 -> Ala** | | | | | | | |
| 7V2C | Std | 0 | + | 2.94 ps | 1[3] | - | def2-SVP |
| 7V2C | Std | 1e- | + | 1.5 ps | 1[3] | - | def2-SVP |
| 7V2C | Std | 2e- | + | 2.92 ps | 1[3] | - | def2-SVP |

[1] "Std" stands for "standard" equilibration protocol, while "Std+Add" introduces an additional 10-ns equilibration stage with constraints on protein backbone, see the "Computational methods" section.

[2] QM region is the same as #1 (see Tabl. S1), but with the mutation of Tyr144 to phenylalanine

[3] QM region is the same as #1 (see Tabl. S1), but with the mutation of His92 to alanine

Table S4. Classical MD simulation setups performed in the present work.

| Protonation states | | | Ubiquinone charge state | Unbiased MD production simulation (time, ns) | | | Umbrella sampling |
|---|---|---|---|---|---|---|---|
| His[1] | Asp | Tyr | | Replica 1 | Replica 2 | Replica 3 | |
| 0 (δ) | 0 | 0 | UQ | ~975 ns | ~972 ns | ~972 ns | + |
| | | | $USQ^-$ | ~966 ns | ~977 ns | ~971 ns | + |
| | | | $UQH^-$ | - | - | - | + |
| +1 | -1 | 0 | UQ | - | - | - | + |
| | | | $USQ^-$ | - | - | - | + |
| | | | $UQH^-$ | - | - | - | + |
| 0 (δ) | 0 | -1 | $UQH^-$ | ~971 ns | ~970 ns | ~962 ns | + |
| 0 (ε) | -1 | 0 | $UQH^-$ | ~1078 ns | ~1095 ns | ~1095 ns | + |
| +1 | -1 | -1 | $UQH^-$ | - | - | - | + |

[1]δ and ε describe the tautomeric state of neutral histidine (proton position on the δ and ε nitrogen, respectively)
